# The time to initiate retrieval of a memory depends on recency

**DOI:** 10.1101/2022.09.16.508287

**Authors:** Ian M. Bright, Inder Singh, Rebecca DiDomenica, Aude Oliva, Marc W. Howard

## Abstract

Continuous recognition has long been used to study the recency effect in recognition memory. In continuous recognition, response time (RT) increases as a logarithmic function of the lag since a probe was last presented. Although this could simply be due to decaying trace strength, careful examination of response time (RT) distributions showed that the onset of RT distributions changed with the logarithm of the lag since a probe was originally presented. Each doubling of lag resulted in a shift of roughly 20 ms in the rise time of the RT distributions. To test the hypothesis that this increase was simply due to increased facility in processing the probe item, Experiment six repeated items six times. Repetition resulted in faster RTs, but did not change the effect of log lag on RT. In light of recent neurophysiological evidence, we consider the hypothesis that memory requires a recovery of temporal context, and the time to retrieve a prior temporal context goes up with the logarithm of the time since it was experienced.

---

In recognition experiments, participants must judge if a given probe was previously experienced or new. As the recency of a repeated probe decreases, the accuracy of this judgment decreases, and response times (RT) increase (Shepard & Teghtsoonian, 1961; Murdock & Anderson, 1975; Monsell, 1978; Hockley, 1982; Donkin & Nosofsky, 2012). Despite decades of empirical and modeling work, the cause of this recency effect remains unclear. In continuous recognition, there is no separation between a study phase and a test phase; the participant must make a new/old judgment on every trial. Consider the task of an individual engaged in continuous recognition. The individual must correctly identify an item as old by comparing it to the contents of their memory. In continuous recognition, the recency effect manifests as a sublinear increase in RT with increasing lag of the repeated probe (Hintzman, 1969; Okada, 1971; Hockley, 1982). Hockley (1982) in particular, found a logarithmic increase in RT with increasing lag. A question that remains however is, why does it take longer to retrieve memories experienced further in the past? One hypothesis is that the strength of memory traces decay over time or with intervening items. A second hypothesis is that it takes longer to retrieve memories from further in the past. These hypotheses are not mutually exclusive.

Strength models are consistent with traditional signal detection models of recognition memory (Murdock & Dufty, 1972; Wixted, 2007; Rotello, 2017), in which the output of memory for each probe consists of a scalar decision variable. Strength models can be implemented in distributed memory models that assume that memory is a composite store containing a noisy record of features from all the studied items (e.g., J. A. Anderson, 1973; Murdock, 1982; Shiffrin, Ratcliff, Murnane, & Nobel, 1993). A composite memory store can account for the recency effect if the features of items experienced further in the past are stored with less fidelity than items experienced more recently. Indeed the confidence/RT relationship (Norman & Wickelgren, 1969; Murdock & Dufty, 1972) is consistent with the hypothesis that old probes that evoke more strength should result in both faster RTs and high confidence in their prior occurrence. A strength model can account for the logarithmic relationship between RT and recency increase if the strength of the match between a probe and the contents of memory decreases appropriately (J. R. Anderson, Bothell, Lebiere, & Matessa, 1998) and if this strength is coupled with a model of information accumulation (e.g., S. Brown & Heathcote, 2005; Ratcliff, 1978; Usher & McClelland, 2001). The key feature of strength models is that information about all traces becomes available at the same time during retrieval.

Another class of models in recognition memory hypothesize that at least some correct judgments in item recognition result from a detailed recollective process (Mandler, 1980; Tulving, 1985; Yonelinas, 1997). Recollection results in the availability of detailed source information about the encoding context of the probe stimulus and this retrieval process can succeed or fail (Province & Rouder, 2012; Kellen & Klauer, 2014). Although the output of recollection need not be constant (Onyper, Zhang, & Howard, 2010) the recollective process should take some time before it provides information about the probe (Hintzman & Curran, 1994). If the time to recover an episodic memory depends on its recency, a discrete state model could also account for the logarithmic effect on RT. It has been proposed that recollection depends on recovery of a gradually-changing temporal context (Tulving & Madigan, 1970; G. Schwartz, Howard, Jing, & Kahana, 2005; Folkerts, Rutishauser, & Howard, 2018). Perhaps the time to retrieve a prior state of temporal context depends on its “distance” from the present state of temporal context. The finding that RT in continuous recognition increases with the log of lag reflects the fact that temporal context changes as a function of log time (G. D. A. Brown, Neath, & Chater, 2007; Howard, Shankar, Aue, & Criss, 2015; Howard, 2018; Cao, Bladon, Charczynski, Hasselmo, & Howard, 2021).

These two hypotheses—the recency effect is due to decaying trace strength *vs* the recency effect is due to a discrete retrieval process that takes an amount of time that depends on recency—can be distinguished by examining the shape of the RT distributions (Figure 1, top). If recency only affects the strength of a memory trace, then the time needed to begin evaluating a given memory is the same regardless of how far in the past the probe was experienced. The strength of that match, however, should depend on the probe’s recency. If, however, old judgments must await the termination of a discrete retrieval process, then RT distributions should rise at different times (Figure 1, bottom). If there is no difference in the quantity of evidence that is retrieved by this discrete retrieval process, then the distributions at different lags will maintain a constant offset from each other as one moves through the distribution. Critically, a change in the time to initiate the search as a function of lag cannot be accounted for by a purely strength based account of the recency effect.

**Figure 1.**
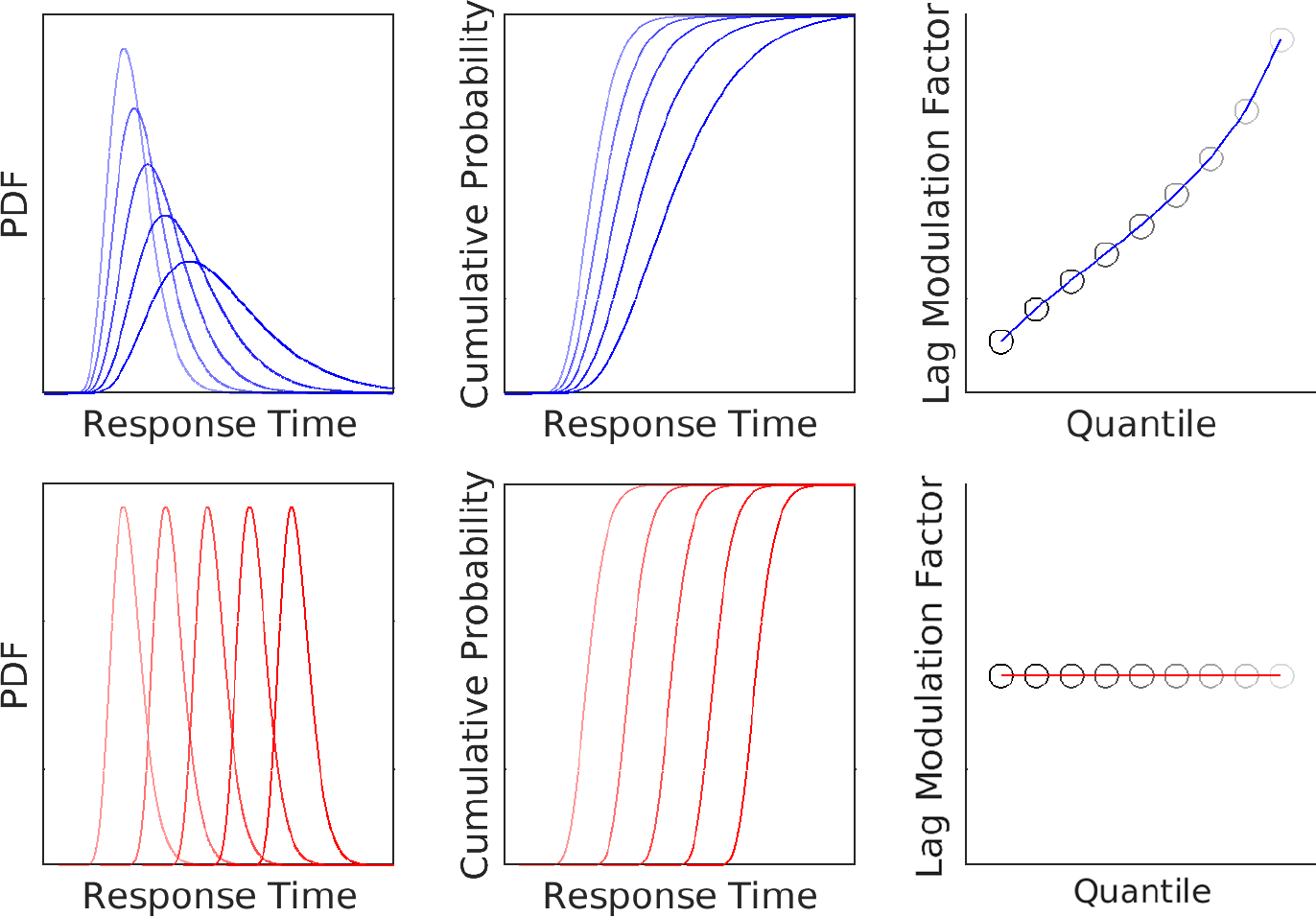
Distinguishing two potential accounts for the recency effect from response time distributions. **Top**. In strength-based accounts of recency, all items are available simultaneously but older items (darker lines) are represented with less fidelity than newer items (lighter lines). *Left*: Probability density functions of response times for each lag. The shape of the distribution changes with the drift rate. *Middle*: A cumulative distribution of response times for different lags. RT distributions start at the same point regardless of lag, but spread as you move into later quantiles. *Right*: A plot of lag modulation factor as a function of quantile. In strength-based accounts of recency, lag modulation factor is zero at the start of the distribution and monotonically increases for later quantiles. **Bottom**. If the recency effect arises due to a delay in recovering older temporal contexts, then older items require more time before evidence accumulation can begin. *Left*: Probability density functions of response times for each lag. Rather than varying in their drift rate, newer items begin accumulating evidence before older items. *Middle*: A cumulative distribution of response times for different lags. Because the distributions do not change their shape, the distance between response times is similar across deciles. *Right*: The Lag Modulation Factor as a function of quantile. The change in the start to accumulate evidence results in a non-zero intercept.

While a shift in the RT distribution is consistent with a recency dependent retrieval process during the memory comparison phase, this is not the only possible explanation. The probe must be encoded before it can be compared to memory. A shift in the RT distributions would also be consistent with the hypothesis that recently-experienced probes are processed faster *prior* to memory retrieval. If the recency of a repeated item allows it to be processed faster as a probe, then repeating the item again should have an additional effect on RT. This implies that the time it takes for an item to become available in memory should depend on the recency of both previous presentations. If changes are driven by improved processing of recently-presented probes, then the recency of both previous presentations ought to both affect RT. In contrast, if the recency effect instead arises due to a recency dependent delay in initiating retrieval of a memory, then the delay would only depend on the most recent lag. Further, this would predict that the effect of recency should be the same on both the first and second presentations.

To the best of our knowledge, a systematic change in the time to initiate the memory search has not previously been observed in continuous recognition. The main issue in continuous recognition is that as lag increases, accuracy decreases (Hockley, 1982; Shepard & Teghtsoonian, 1961), making it more challenging to measure the effect of recency on retrieval dynamics independently of changes in accuracy. Brady, Konkle, Alvarez, and Oliva (2008) showed participants hundreds of memorable images in a continuous recognition task with lags varying over more than two orders of magnitude, with lags from 1 (no intervening items) to 128. In addition, the RT data from Brady et al. (2008) are minimally affected by sequential dependencies, which are known to affect RTs in recognition memory (Malmberg & Annis, 2012), as repeated items could not take place in adjacent trials. Because of the wide range of lags tested, and the elimination of sequential dependencies, Brady et al. (2008) is well-suited to study the effect of recency on RT distributions. This paper analyzes the RT data collected during the Brady et al. (2008) task (referred to as Experiment 1), and five additional experiments designed to assess the generality of this finding. All six experiments show a systematic change in the rise time of RT distributions as a function of log lag, suggesting a discrete retrieval process that depends on the recency of the to-be-retrieved memory. Experiment 6 assesses the alternative hypothesis that the change in the rise time of RT distributions is due to probe fluency by repeating old items up to five times, presumably saturating probe fluency. Despite an overall decrease in RT for repeated items—indicating that the manipulation of probe fluency was successful—the effect of recency on RT was unchanged.

## Methods

Table 1 contains a summary of the major methodological differences between the six continuous recognition experiments. In short, experiment 2 was a replication of experiment 1 (Brady et al., 2008) using a different subject population (i.e., Amazon M-Turk or Boston University Students). Experiments 3-6 displayed phase scrambled noise rather than a fixation cross between each image. In experiment 4, words from the Toronto Word Pool (Friendly, Franklin, Hoffman, & Rubin, 1982) were used as stimuli rather than images. In experiments 1-4, participants were only required to respond if they believed the stimuli had previously been presented, in experiments 5-6 participants had to respond new/old on every trial. Finally in experiments 1-5, some stimuli were repeated a second time while in experiment 6 some stimuli were repeated five times.

**Table 1:**
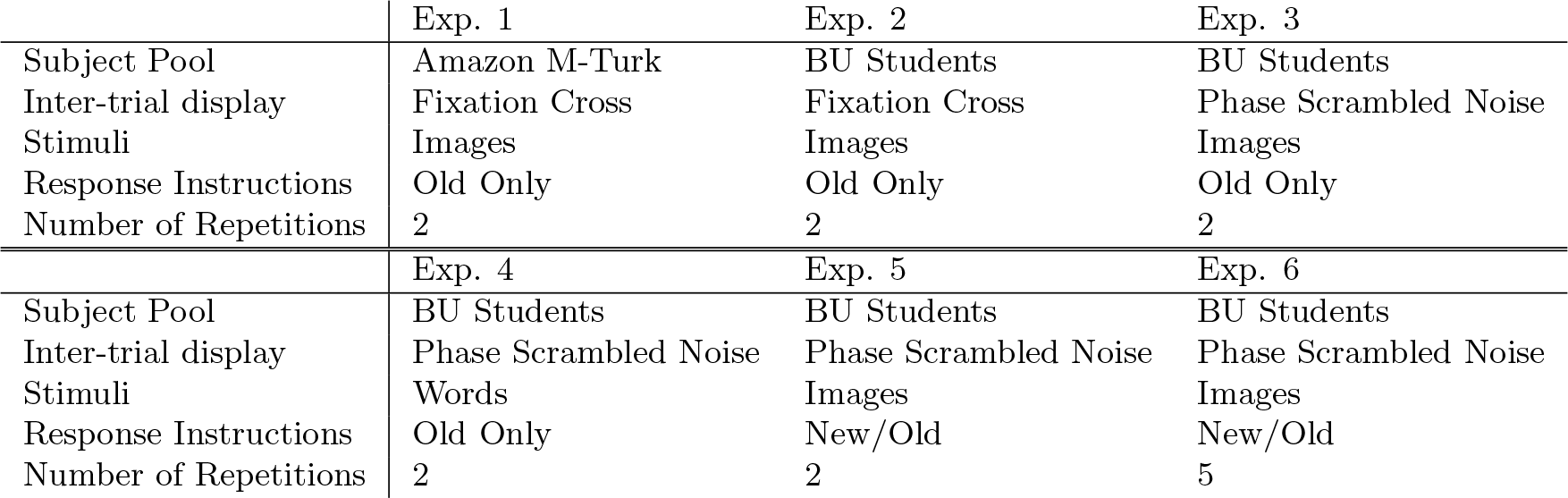
The major methodological differences between the six continuous recognition experiments. The experiments varied on the subject pool that participants were recruited from (Amazon M-Turk or Boston University Students), what was displayed between each trial (fixation cross or phase scrambled noise), the type of stimuli used (images or words), the instructions given to the participants (respond only to repeated items or indicate if the current item is new/old), and the number of times an item could be repeated (2 or 5).

### Experiment 1

#### Participants

Fourteen individuals participated in this study in exchange for financial compensation. Participants were recruited via Amazon Mechanical Turk. All participants completed the task simultaneously, and completed the task at computer workstations that were matched for similar screen size and viewing distance.

#### Materials

Stimuli were assembled from a commercially available database (Hemera Photo-Objects, Vol. I and II) and searches done on Google Image Search. Each participant viewed a series of 2500 images, 396 of which were randomly chosen to be repeated. Of these 396 images, 56 were repeated immediately (lag 1), 52 were repeated such that the previous presentation was two trials prior (lag 2), 48 were repeated four trials after their previous presentation, and so on out to 16 images repeated at lag 1024. Lags greater than 128 were not considered, as repetitions occurred across blocks. No two consecutive trials were allowed to contain repeated images, preventing sequential effects. Some images were repeated twice, at lags greater than 1, to avoid adjacent repeat trials. In order to minimize the effect of fatigue, and to simplify counterbalancing when repeats of various lags would occur, the trials with repeated items were the same for all participants. However, all participants viewed images in different orders, and different images were repeated for each participant.

#### Procedure

Participants viewed ten blocks of images, which lasted around 20 minutes each. All blocks contained 290 trials, except for the first block which consisted of 289 trials, and the tenth and final block which consisted of 287 trials. Participants were given a 5 minute break between each block, and instructed to not discuss any images they had seen. Images were presented for 3 seconds, with a fixation cross presented for 800 ms between each image. The stimuli, approximately 7.5° by 7.5° of visual angle, were presented one at a time in the center of the screen. Participants were instructed to hit the space bar if the current image had previously been presented. They did not need to do anything on trials where they judged the current image to be new. Participants only received feedback on trials in which they responded. On false alarm trials the fixation cross following the trial was red, and on trials which were hits, the fixation cross turned green.

### Experiment 2

#### Participants

Twenty-nine individuals participated and were compensated with $15 for their time. All participants were members of the Boston University community, and were recruited via the quick job page. Participants gave informed written consent prior to beginning the study, and the protocol was approved by the Boston University Charles River Institutional Review Board. Participants completed the task at different times at one of two 21.5 inch, mid-2011 iMac computers.

#### Materials

The experiment was implemented in PyEPL (Geller, Schlefer, Sederberg, Jacobs, & Kahana, 2007). Stimuli were pulled from the same set of images as used in experiment 1. Each participant viewed a series of 650 images, 250 of which were randomly chosen to be repeated. Of those 250 images, 30 were repeated immediately (lag 1), 28 were repeated at a lag of 2, and so on out to 16 images repeated at lag 128. Repetitions that occurred across blocks were not considered. For each subject, 27 items were repeated a second time at lags 2, 4, and 16 (9 per lag). For items repeated a second time, the lag between its first presentation and first repetition was restricted to 8, 16, and 64. The presentation order of images and the trials which contained repetitions were randomized for each subject.

#### Procedure

Participants viewed two blocks of images, which lasted around 20 minutes each. Both blocks consisted of 450 trials. Participants were given a short break (5 minutes) between the two blocks. Images were presented for 2.6 seconds, with a fixation cross presented for 400 ms between each image. The stimuli, approximately 7.5° by 7.5° of visual angle, were presented one at a time in the center of the screen. Participants were instructed to hit the space bar if the current image had previously been presented. They did not need to respond on trials where they judged the current image to be new. Participants only received feedback on trials in which they responded. On false alarm trials the fixation cross following the trial was red, and on trials which were hits, the fixation cross turned green.

### Experiment 3

#### Participants

Thirty-nine individuals participated and were compensated with $15 for their time. All participants were members of the Boston University community, and were recruited via the quick job page. Participants gave informed written consent prior to beginning the study, and the protocol was approved by the Boston University Charles River Institutional Review Board. Participants completed the task at different times at one of two 21.5 inch, mid-2011 iMac computers.

#### Materials

The experiment was implemented in PyEPL (Geller et al., 2007). Stimuli were assembled from the same set of images as used in the previous experiments. In this experiment, there was a wider range of lag values, ranging from lag 1 to lag 512. In order to better accommodate these longer lags, participants viewed a series of 1360 images, 276 of which were randomly chosen to be repeated. Of those 276 repeated images, 30 were chosen to be repeated immediately (lag 1), 28 were repeated at a lag of 2, and so on out to 12 images repeated at lag 512. Repetitions that occurred across blocks were not considered. For each subject, 27 items were repeated a second time at lags 2, 4, and 16 (9 per lag). For items repeated a second time, the lag between its first presentation and first repetition was restricted to 8, 16, and 64. As in experiment 2, both the order of the images, and the trials where repeated items occurred were randomized for each participant.

#### Procedure

Participants viewed two blocks of images, which lasted around 35 minutes each. Both blocks consisted of 680 trials. Participants were given a short break (5 minutes) between the two blocks. Images were presented for 2.6 seconds, however instead of a fixation cross, phase scrambled images were used as masked separators and presented for 400 ms between each image. The stimuli, approximately 7.5° by 7.5° in visual angle, were presented one at a time in the center of the screen. Participants were instructed to hit the space bar if the current image had previously been presented. They did not need to do anything on trials where they judged the current image to be new. Participants only received feedback on trials in which they responded. On false alarm trials a red square appeared around the image and on trials which were hits, a green square appeared around the image.

### Experiment 4

#### Participants

Thirty-three individuals participated and were compensated with $15 for their time. All participants were members of the Boston University community, and were recruited via the quick job page. Participants gave informed written consent prior to beginning the study, and the protocol was approved by the Boston University Charles River Institutional Review Board. Participants completed the task at different times at one of two 21.5 inch, mid-2011 iMac computers.

#### Materials

The experiment was implemented in PyEPL (Geller et al., 2007). Stimuli were assembled from the Toronto Word Pool (Friendly et al., 1982), a list of common English nouns. Participants viewed a series of 1360 words, 276 of which were randomly chosen to be repeated. Of those 276 repeated words, 30 were chosen to be repeated immediately (lag 1), 28 were repeated at a lag of 2, and so on out to 12 images repeated at lag 512. Repetitions that occurred across blocks were not considered. For each subject, 27 items were repeated a second time at lags 2, 4, and 16 (9 per lag). For items repeated a second time, the lag between its first presentation and first repetition was restricted to 8, 16, 64. Both the order of the words, and the trials containing repeat items were randomized for each participant.

#### Procedure

Participants viewed two blocks of words, which lasted around 35 minutes each. Both blocks consisted of 1360 trials. Participants were given a short break (5 minutes) between the two blocks. As in experiment 3, stimuli were presented for 2.6 seconds, and phase scrambled images were used as masked separators and presented for 400 ms between each image. The stimuli, approximately 7.5° by 7.5° of visual angle, were presented one at a time in the center of the screen. Participants were instructed to hit the space bar if the current word had previously been presented. They did not need to do anything on trials where they judged the current word to be new. Participants only received feedback on trials in which they responded. On false alarm trials a red square appeared around the word and on trials which were hits, a green square appeared around the word.

### Experiment 5

#### Participants

Fourty-one individuals participated and were compensated with $15 for their time. All participants were members of the Boston University community, and were recruited via the quick job page. Participants gave informed written consent prior to beginning the study, and the protocol was approved by the Boston University Charles River Institutional Review Board. Participants completed the task at different times at one of two 21.5 inch, mid-2011 iMac computers.

#### Materials

Stimuli were assembled from the same set of images as used in experiments 1-3. Participants viewed a series of 1360 images, 276 of which were randomly chosen to be repeated. Of those 276 repeated images, 30 were chosen to be repeated immediately (lag 1), 28 were repeated at a lag of 2, and so on out to 12 images repeated at lag 512. Repetitions that occurred across blocks were not considered. For each subject, 27 items were repeated a second time at lags 2, 4, and 16 (9 per lag). For items repeated a second time, the lag between its first presentation and first repetition was restricted to 8, 16, and 64. Both the order of the images, and the trials where repeated items occurred were randomized for each participant.

#### Procedure

Participants viewed two blocks of words, which lasted around 35 minutes each. Both blocks consisted of 680 trials. Participants were given a short break (5 minutes) between the two blocks. As in experiment 3, stimuli were presented for 2.6 seconds, and phase scrambled images were used as masked separators and presented for 400 ms between each image. The stimuli, approximately 7.5° by 7.5° of visual angle, were presented one at a time in the center of the screen. Different from previous experiments, participants were instructed to hit the left arrow key if the current image was repeated, and the right arrow key if the current image was new. Participants only received feedback on trials in which they responded. On incorrect trials (false alarms and misses) a red square appeared around the image and on correct trials (hits and correct rejections) a green square appeared around the image.

### Experiment 6

#### Participants

Fourty-six individuals participated and were compensated with $15 for their time. All participants were members of the Boston University community, and were recruited via the quick job page. Participants gave informed written consent prior to beginning the study, and the protocol was approved by the Boston University Charles River Institutional Review Board. Participants completed the task at different times at one of two 21.5 inch, mid-2011 iMac computers.

#### Materials

Stimuli were assembled from the same set of images as used in experiment 5. Participants viewed a series of 1360 images, 276 of which were randomly chosen to be repeated. Of those 276 repeated images, 30 were chosen to be repeated immediately (lag 1), 28 were repeated at a lag of 2, and so on out to 12 images repeated at lag 512. Repetitions that occurred across blocks were not considered. For each subject, 27 items were repeated five times. For items repeated more than once, the lag between its first presentation and first repetition was restricted to 8, 16, and 64. For all subsequent repetitions, the possible lags were 2, 4, and 16 (9 per lag and number of repetitions). Both the order of the images, and the trials where repeated items occurred were randomized for each participant.

#### Procedure

Participants viewed two blocks of words, which lasted around 35 minutes each. Both blocks consisted of 1360 trials. Participants were given a short break (5 minutes) between the two blocks. Stimuli were presented for 2.6 seconds, and phase scrambled images were used as masked separators and presented for 400 ms between each image. The stimuli, approximately 7.5° by 7.5° of visual angle, were presented one at a time in the center of the screen. As in experiment 5, participants were instructed to hit the left arrow key if the current image was repeated, and the right arrow key if the current image was new. Participants only received feedback on trials in which they responded. On incorrect trials (false alarms and misses) a red square appeared around the image and on correct trials a green square appeared around the image.

#### Analyses

Analyses were performed using R statistical software. Both linear and logistic regressions were performed as mixed effects regressions, using the NLME package in R (Pinheiro, Bates, DebRoy, Sarkar, & R Core Team, 2021). All analyses were performed on the base 2 log of the lag. Due to the possibility that responses for items repeated immediately may not necessarily involve retrieval from memory, lag 1 repetitions were not considered in any regressions. Lags larger than 128 were also not considered. Within Subject error bars were calculated using the method outlined in Morey (2008).

All data and code used to perform analyses have been made publicly available on Github and can be accessed at https://github.com/tcnlab/ConRec.

## Results

To anticipate the results, the data from all six experiments showed evidence that the time to retrieve a memory, as operationalized by the onset time of the RT distribution, changed systematically with the recency of the probe item. When considering probes repeated once, they were faster and more likely to be recognized the more recently an item was last presented. Analysis of response time distributions showed an effect of recency even for the fastest response times. Analysis of multiple repetitions found that correct RT’s decreased with the number of previous presentations. Critically, the time to access a previously presented probe only depended on the recency of its most recent presentation. Further, the effect of recency was the same regardless of the number of repetitions. In all figures, experiment 1 corresponds to black circles and solid lines, experiment 2 corresponds to black triangles and dashed lines, experiment 3 corresponds to blue circles and solid lines, experiment 4 corresponds to blue triangles and dashed lines, experiment 5 corresponds to red circles and solid lines, and experiment 6 corresponds to red triangles and dashed lines.

### First Repetitions

#### Items presented more recently were more likely to be recognized

The average false alarm rate was low across experiments. In experiment 1, the false alarm rate was 1.4 percent, the highest hit rate was at lag 1 at 99.6 percent (corresponding to a d-prime of 4.85) and lowest at lag 128 at 89.7 percent (*d′* = 3.46). In experiment 2, the false alarm rate remained low across participants at 3.8 percent, hit rate was highest for lag 1 at 96.2 percent (*d′* = 3.55), and lowest for lag 128 with an average hit rate of 69.2 percent (*d′* = 2.27). Experiment 3 was similar to experiment 2, the false alarm rate was 3.0 percent, hit rate was the highest for lag 1 at 91.9 percent (*d′* = 3.28), and lowest for lag 128 with an average hit rate of 69.2 percent (*d′* = 3.28). In experiment 4, which used words as stimuli, the false alarm rate was higher than in previous experiments at 6.9 percent, hit rate was highest for lag 1 at 94.2 percent (*d′* = 3.06), and lowest for lag 128 with an average hit rate of 61.5 percent (*d′* = 1.78). In experiment 5, which used the same stimuli as experiments 1-3 but required a new/old response, the false alarm rate was similar to those experiments at 2.0 percent, but overall hit rate was lower. Hit rate was highest for lag 1 at 86.1 percent (*d′* = 3.12) and lowest for lag 128 with an average hit rate of 46.2 percent (*d′* = 1.94). Experiment 6, which also required new/old responses for each trial was resulted in a similar false alarm rate of 2.6 percent, hit rate was highest for lag 1 at 84.4 percent (*d′* = 2.96), and lowest for lag 128 with an average hit rate of 46.2 percent (*d′* = 2.03). Subjects were able to distinguish old probes from new probes successfully out to at least six minutes.

Average subject hit rate as a function of log lag is plotted in Figure 2a. There was a recency effect in accuracy for all experiments. In order to quantify the decrease in accuracy for increasing lags, we performed a mixed effects logistic regression on hit rate, treating log lag as a fixed effect and subject as a random effect. Table 2 shows the results of this analysis for each experiment. Across all six experiments, hit rate significantly decreased with each doubling of lag (*p* < 0.001). Subjects we less likely to correctly recognize repeated items the further in the past they were presented.

**Figure 2.**
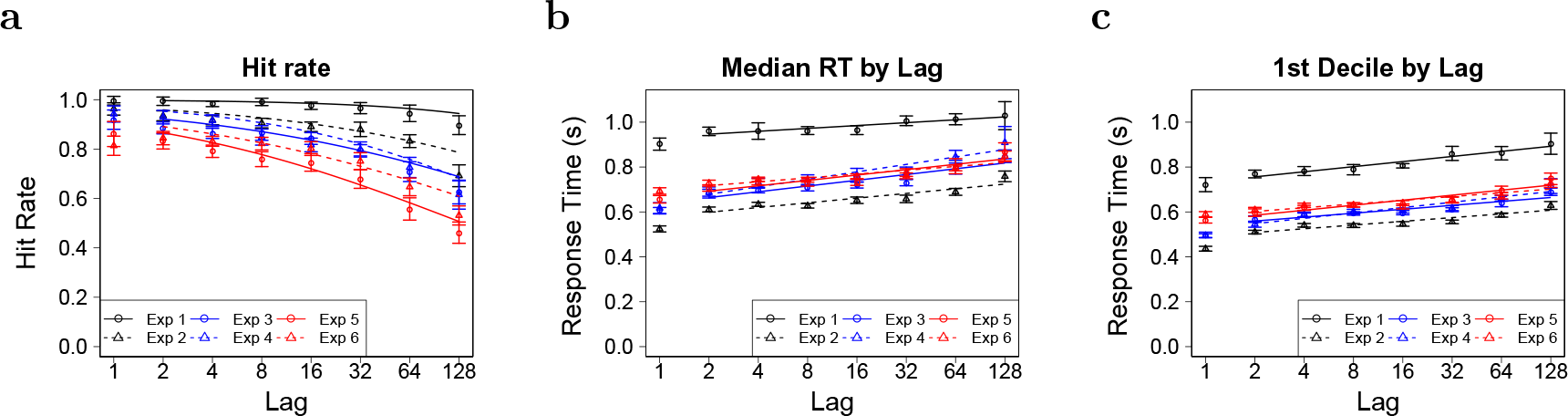
The recency effect is robust across experimental conditions. **a**. Hit Rate as a function of log_2_ lag for Experiments 1-6. The hit rate goes down with lag. There is a more pronounced drop in the hit rate at higher lags. **b**. Median response time as a function of log2 lag. Median response time increased linearly with the logarithm of lag. **c**. The 1st decile as a function of log2 lag. 1st decile response times increased linearly with the logarithm of the lag at a similar rate as median response times. Error bars in all figures represent the 95% confidence interval of the mean across participants normalized using the method described in Morey (2008).

**Table 2:** Slope and intercept values from a logistic mixed effects regression of lag on hit rate for first repetition items. All tests were performed on the base 2 logarithm of lag. Bold z-scores indicate significance at the *p* < 0.05 level. Across all six experiments, hit rates decreased as a function of lag.

| Hit Rate | Exp. 1 | Exp. 2 | Exp. 3 |
| --- | --- | --- | --- |
| Intercept | $6.04 \pm 0.47$<br>$z = \mathbf{12.75}, p < 0.001$ | $3.45 \pm 0.18$<br>$z = \mathbf{19.18}, p < 0.001$ | $2.74 \pm 0.16$<br>$z = \mathbf{17.54}, p < 0.001$ |
| Slope | $-0.46 \pm 0.07$<br>$z = \mathbf{-6.72}, p < 0.001$ | $-0.31 \pm 0.02$<br>$z = \mathbf{-13.88}, p < 0.001$ | $-0.28 \pm 0.02$<br>$z = \mathbf{-15.27}, p < 0.001$ |
|  | Exp. 4 | Exp. 5 | Exp. 6 |
| Intercept | $3.39 \pm 0.22$<br>$z = \mathbf{15.17}, p < 0.001$ | $2.17 \pm 0.15$<br>$z = \mathbf{14.59}, p < 0.001$ | $2.38 \pm 0.15$<br>$z = \mathbf{16.22}, p < 0.001$ |
| Slope | $-0.37 \pm 0.02$<br>$z = \mathbf{-17.53}, p < 0.001$ | $-0.31 \pm 0.02$<br>$z = \mathbf{-16.90}, p < 0.001$ | $-0.28 \pm 0.02$<br>$z = \mathbf{-15.85}, p < 0.001$ |

**Table 3:** Slope and intercept values from a linear mixed effects regression of lag on median response times for first repetition items. All tests were performed on the base 2 logarithm of lag. Bold t-scores indicate significance at the *p* < 0.05 level. Across all six experiments, response times increased as a function of lag.

| Median | Exp. 1 | Exp. 2 | Exp. 3 |
| --- | --- | --- | --- |
| Intercept | $0.934 \pm 0.042$<br>$t(83) = \mathbf{22.22}, p < 0.001$ | $0.578 \pm 0.013$<br>$t(221) = \mathbf{45.05}, p < 0.001$ | $0.639 \pm 0.021$<br>$t(227) = \mathbf{30.16}, p < 0.001$ |
| Slope | $0.013 \pm 0.003$<br>$t(83) = \mathbf{4.54}, p < 0.001$ | $0.021 \pm 0.002$<br>$t(221) = \mathbf{13.65}, p < 0.001$ | $0.026 \pm 0.002$<br>$t(227) = \mathbf{13.20}, p < 0.001$ |
|  | Exp. 4 | Exp. 5 | Exp. 6 |
| Intercept | $0.646 \pm 0.022$<br>$t(197) = \mathbf{28.88}, p < 0.001$ | $0.669 \pm 0.020$<br>$t(179) = \mathbf{32.85}, p < 0.001$ | $0.700 \pm 0.016$<br>$t(209) = \mathbf{42.55}, p < 0.001$ |
| Slope | $0.033 \pm 0.003$<br>$t(197) = \mathbf{9.69}, p < 0.001$ | $0.024 \pm 0.002$<br>$t(179) = \mathbf{10.52}, p < 0.001$ | $0.017 \pm 0.001$<br>$t(209) = \mathbf{11.98}, p < 0.001$ |

#### More recent items were recognized faster than older items

Figure 2b shows the average median correct response time across subjects as a function of log lag. There was a recency effect on median response times; more recently presented items were recognized faster than less recent items. To confirm the existence of this recency effect, a mixed effects linear regression, treating subject as a random effect and log lag as a fixed effect was performed. Table 4 shows the results of this analysis for each experiment. Across all six experiments, median response times significantly increased for each doubling of lag (*p* < 0.001). Subjects recognized items faster the more recently they were presented.

**Table 4:** Slope and intercept values from a linear mixed effects regression of lag on first decile response times for first repetition items. All tests were performed on the base 2 logarithm of lag. Bold t-scores indicate significance at the *p* < 0.05 level. Across all six experiments, response times increased as a function of lag.

| 1st Decile | Exp. 1 | Exp. 2 | Exp. 3 |
| --- | --- | --- | --- |
| Intercept | 0.734 ± 0.033<br>$t(83) = \mathbf{22.05}, p < 0.001$ | 0.492 ± 0.008<br>$t(221) = \mathbf{60.63}, p < 0.001$ | 0.542 ± 0.012<br>$t(227) = \mathbf{45.17}, p < 0.001$ |
| Slope | 0.023 ± 0.002<br>$t(83) = \mathbf{9.29}, p < 0.001$ | 0.017 ± 0.001<br>$t(221) = \mathbf{15.65}, p < 0.001$ | 0.018 ± 0.001<br>$t(227) = \mathbf{14.22}, p < 0.001$ |
|  | Exp. 4 | Exp. 5 | Exp. 6 |
| Intercept | 0.524 ± 0.012<br>$t(197) = \mathbf{41.91}, p < 0.001$ | 0.563 ± 0.014<br>$t(179) = \mathbf{39.26}, p < 0.001$ | 0.586 ± 0.013<br>$t(209) = \mathbf{44.59}, p < 0.001$ |
| Slope | 0.024 ± 0.002<br>$t(197) = \mathbf{14.76}, p < 0.001$ | 0.022 ± 0.002<br>$t(179) = \mathbf{13.66}, p < 0.001$ | 0.017 ± 0.001<br>$t(209) = \mathbf{13.23}, p < 0.001$ |

As shown in Figure 1, the two classes of models that explain the recency effect make differing predictions about how the leading edge of their distributions should vary with lag. Figure 3 shows the empirical cumulative response time distributions for all lags. The curves rise at different points, with smaller lags rising earlier than larger lags, indicating an effect of lag for even the fastest responses. As leading edge of the distribution is well described by the first decile (Ratcliff & Smith, 2004; Smith & Ratcliff, 2009), evidence of a recency effect at this point in the distribution would indicate a shift in the RT distributions. Figure 2c shows the average first decile correct response time across subjects as a function of log lag. There was a recency effect present for the fastest responses. A mixed effects linear regression on 1st decile response times, treating subject as a random effect and log lag as a fixed effect confirmed this visual impression. Table 4 shows the results of this analysis for each experiment. For each doubling of lag, first decile response times significantly increased. These findings demonstrate that there was a recency effect for even fastest response times, consistent with the shift predicted by a recency dependent retrieval process.

**Figure 3.**
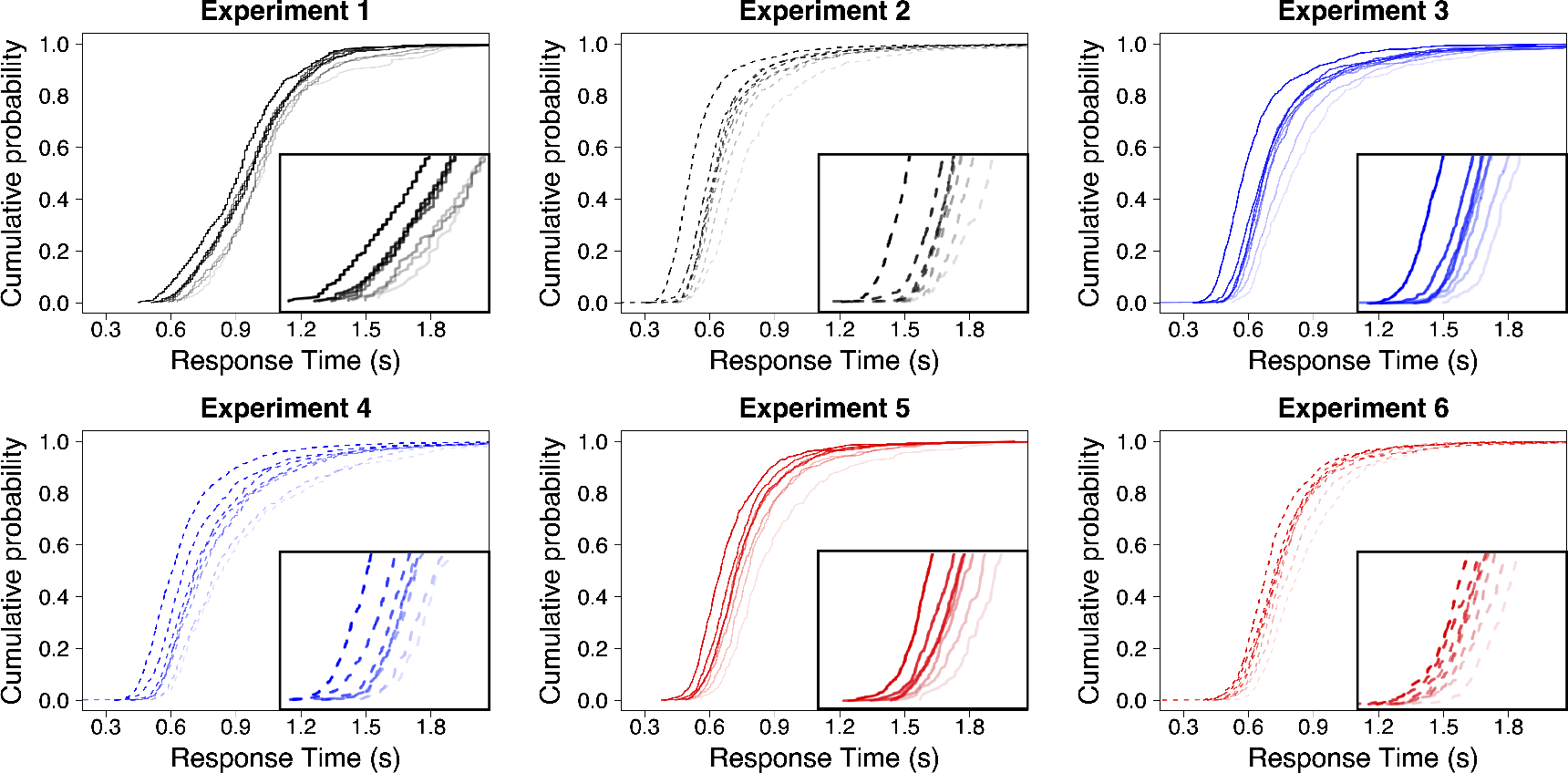
Time to access memory changed systematically with lag in all six experiments. Unsmoothed across-participant cumulative response distributions for each lag. Shorter lags correspond to darker lines. Note that the cumulative distributions shift with decreasing recency. The inset consists of a zoomed in view of the rising points of the distributions. For all six experiments, there is consistent evidence that newer items are available before older items.

#### Non-parametric analyses of response time distributions indicate that more recent memories are available earlier than less recent memories

In order to better formalize this shift, particularly for the fastest responses, we calculated the slope of each subject’s response times as a function of log lag, which we refer to as its “Lag Modulation Factor”, and measured it at different quantiles. As illustrated in Figure 1, a strength-based account predicts that Lag Modulation Factor is zero for the fastest responses, and increases later in the distribution. If instead, recency determines when a memory can be compared to the probe, then Lag Modulation Factor is non-zero, even for the fastest responses. The results of the Lag Modulation Factor analysis offered substantial support to this view. For all six experiments, a mixed effects linear regression of lag modulation factor onto quantile, treating subject as a random effect and quantile as a fixed effect was fit to the data. Table 5 shows the results of this analysis for the six experiments. Critically, in all six experiments lag modulation factor was significant at the intercept (*p* < 0.001), indicating there was an effect of lag on response time for even the fastest responses. The rate at which lag modulation factor changed throughout the distribution varied across experiments, casting further doubts on the ability of a composite model to account for this pattern of results. In experiments 2 (*p* < 0.01), 3 and 4 (*p* < 0.001), lag modulation factor significantly increased, consistent with a drift rate that is slower for less recent items. This is complicated however by the finding that lag modulation factor did not significantly change with quantile in experiments 5 and 6 (*p* > 0.1), and significantly decreased in experiment 1 (*p* < 0.001). This pattern of results is also striking when looking at the subject level. Figure 4 plots the intercept values for each subject’s lag modulation factor as calculated by a linear regression of Lag Modulation Factor onto quantile. With the exception of a single subject in experiments 2 and 3, all participants in all experiments had a positive lag modulation factor at the intercept. Taken together, these results provide strong support that the recency effect in continuous recognition is driven by a change in the time at which items become available.

**Table 5:**
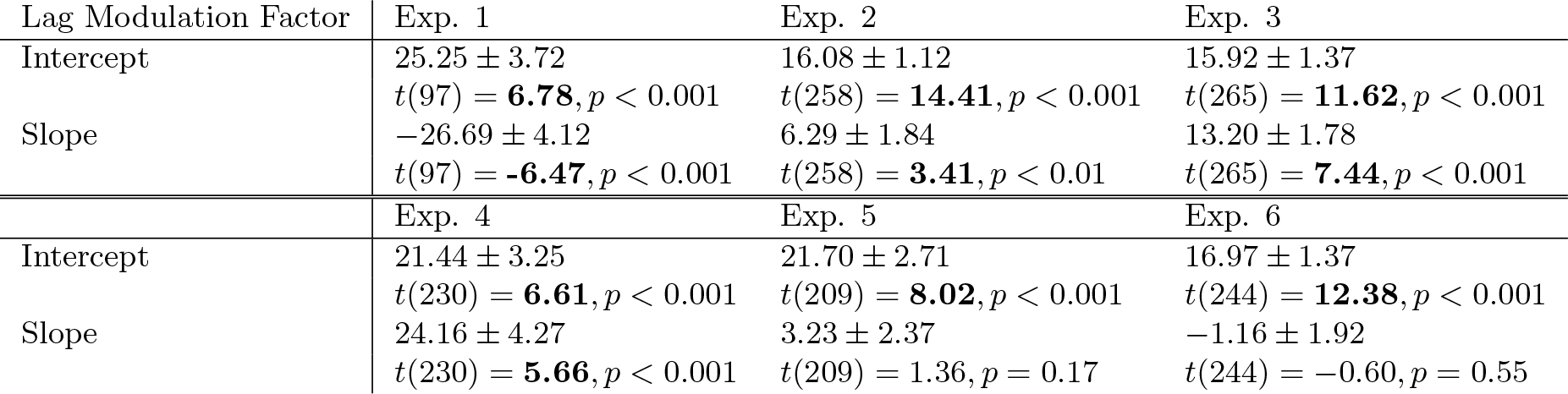
The slopes and intercepts of the linear mixed effects regression on Lag Modulation Factor as a function of Quantile across the six experiments for first repetition items. Bold t-scores indicate significance at the *p* < 0.05 level. The intercepts calculated from the lag modulation factor analysis were significant in all six experiments.

| Lag Modulation Factor | Exp. 1 | Exp. 2 | Exp. 3 |
| --- | --- | --- | --- |
| Intercept | 25.25 ± 3.72 | 16.08 ± 1.12 | 15.92 ± 1.37 |
| | $t(97) = \mathbf{6.78}, p < 0.001$ | $t(258) = \mathbf{14.41}, p < 0.001$ | $t(265) = \mathbf{11.62}, p < 0.001$ |
| Slope | -26.69 ± 4.12 | 6.29 ± 1.84 | 13.20 ± 1.78 |
| | $t(97) = \mathbf{-6.47}, p < 0.001$ | $t(258) = \mathbf{3.41}, p < 0.01$ | $t(265) = \mathbf{7.44}, p < 0.001$ |
|  | Exp. 4 | Exp. 5 | Exp. 6 |
| Intercept | 21.44 ± 3.25 | 21.70 ± 2.71 | 16.97 ± 1.37 |
| | $t(230) = \mathbf{6.61}, p < 0.001$ | $t(209) = \mathbf{8.02}, p < 0.001$ | $t(244) = \mathbf{12.38}, p < 0.001$ |
| Slope | 24.16 ± 4.27 | 3.23 ± 2.37 | -1.16 ± 1.92 |
| | $t(230) = \mathbf{5.66}, p < 0.001$ | $t(209) = 1.36, p = 0.17$ | $t(244) = -0.60, p = 0.55$ |

**Figure 4.**
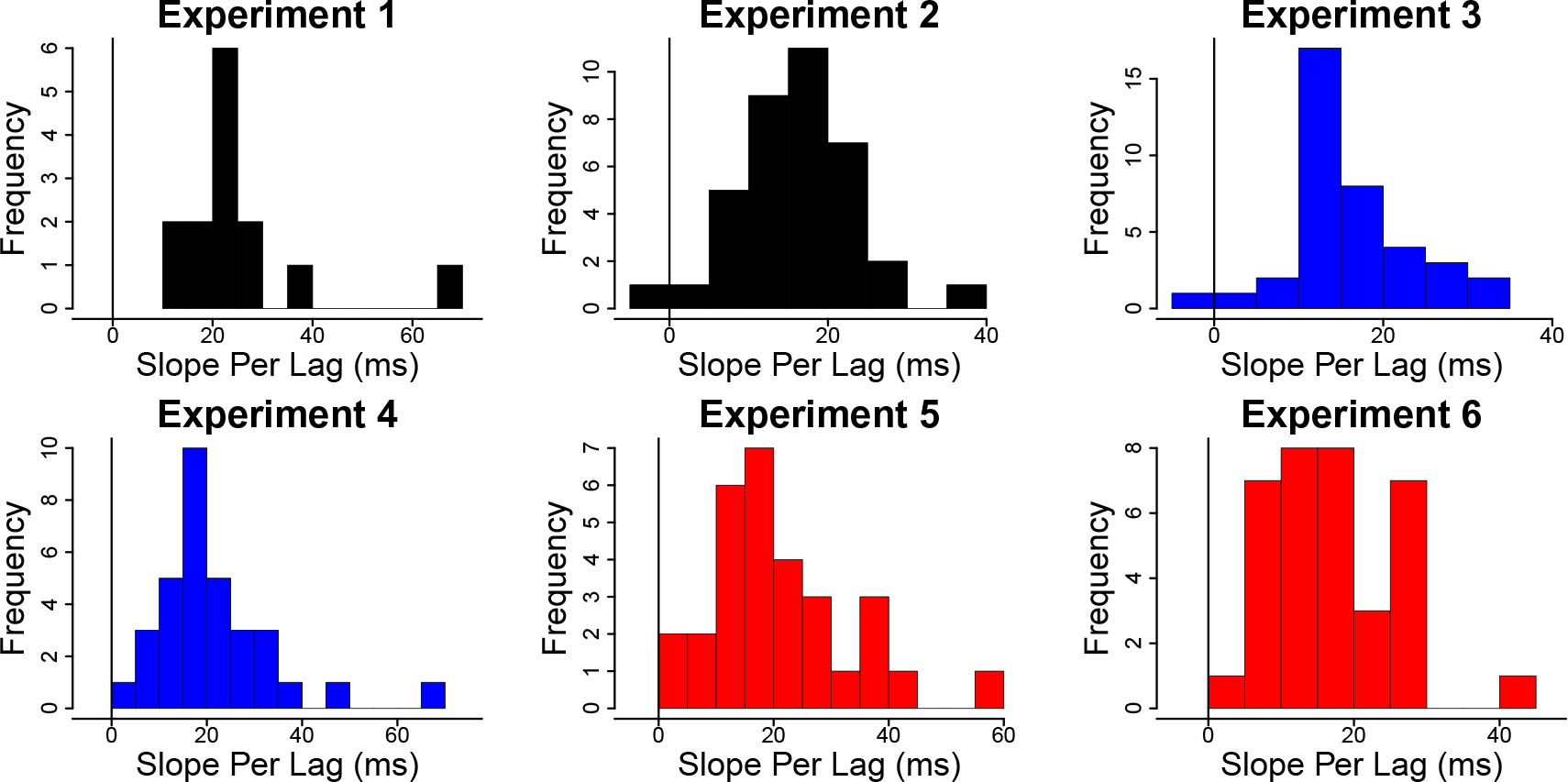
Recency impacted response times for even the fastest responses. Histograms of each subject’s slope per lag (ms) for the fastest responses as calculated by the Lag Modulation Factor analysis. Across all six experiments, subjects had significant recency effects at the start of their RT distributions.

### Multiple Repetitions

Analysis of first repetition responses showed evidence that recency changes response time distributions via a shift in the distribution, consistent with a change in the time to access memory. In order to consider the hypothesis that the recency effect was attributable to changes in encoding, we examined response times for probes repeated more than once.

#### Only the most recent lag impacted response times for items repeated multiple times

Response times depended only on the most recent lag. We performed a mixed effects linear regression of median response times for items repeated a second time, treating the most recent lag (lag_2_) and the lag between the first and second presentations of the image (lag_1_) as fixed effects and subject as a random effect (Table 6). In experiments 1-5, this analysis found that doubling lag_2_ significantly increased median response times, while doubling lag_1_ did not have a significant effect. In experiment 6, while doubling lag_2_ resulted in a significant increase in response times, unlike in previous experiments, doubling lag_1_ also resulted in a significant increasing in response times of. Subsequent analyses suggest this divergence in results was an artifact of how the lists were assembled. In experiment 6, which was kept consistent with experiments 2-5, lag_1_ was confounded with number of repetitions. For items repeated a second time, lag_1_ could equal 8, 16, and 64, for all subsequent repetitions, lag_1_ could equal 2, 4, and 16. It is possible that response times do not increase with lag_1_, but instead our analysis is capturing that second repetition items are slower than items repeated 3 or more times. In order to determine if this was the case, two follow up analyses were performed. When considering only items repeated three or more times, removing the confound of lag_1_ and repetition, only the most recent lag impacted response times. As before, we performed a mixed effects linear regression of median response times for items repeated a three or more times, treating the most recent lag (lag_2_) and the lag between the previous repetition and its previous presentation (lag_1_) as fixed effects and subject as a random effect. We found a significant effect of lag_2_ on RT (0.011 ± 0.002, *t*(278) = 6.77, *p* < 0.001), but no significant effect of lag_1_ (−0.001 ± 0.002, *t*(488) = −0.31, *p* = 0.75). As an additional control, we performed the same mixed effects linear regression for median response times on items repeated twice or more as before, but included a fixed variable that was 1 if a trial contained the second repetition of an item, and zero for all other repetitions in order to separate the effect of (lag_1_) from a repetition effect. We found significant effects of lag_2_ (0.008 ± 0.002, *t*(592) = 4.25, *p* < 0.001) and repetition (0.058 ± 0.006, *t*(592) = 9.30, *p* < 0.001), but lag_1_ did not significantly effect response times (−0.002 ± 0.002, *t*(592) = −1.36, *p* = 0.17). These subsequent analyses suggest that our results in experiment 6 are in line with the previous experiments, and indicate that the recency effect is not the result of improved encoding for more recently presented items.

**Table 6:** Slope and intercept values from a linear mixed effects regression of the lag between an item’s second and third presentation (lag_2_) and the lag between an item’s first and second presentation (lag_1_) and on median response times for items repeated a second time. All tests were performed on the base 2 logarithm of lag. Bold t-scores indicate significance at the *p* < 0.05 level. Across all six experiments, response times increased as a function of lag_2_ but did not systematically vary for lag_1_.

| Median | Exp. 1 | Exp. 2 | Exp. 3 |
| --- | --- | --- | --- |
| Intercept | $0.871 \pm 0.041$<br>$t(393) = \mathbf{21.11}, p < 0.001$ | $0.534 \pm 0.015$<br>$t(294) = \mathbf{35.71}, p < 0.001$ | $0.605 \pm 0.022$<br>$t(302) = \mathbf{27.04}, p < 0.001$ |
| Lag 2 | $0.021 \pm 0.005$<br>$t(393) = \mathbf{4.34}, p < 0.001$ | $0.012 \pm 0.002$<br>$t(294) = \mathbf{5.37}, p < 0.001$ | $0.012 \pm 0.003$<br>$t(302) = \mathbf{4.43}, p < 0.001$ |
| Lag 1 | $-0.004 \pm 0.004$<br>$t(393) = -0.83, p = 0.41$ | $-0.001 \pm 0.002$<br>$t(294) = -0.54, p = 0.59$ | $-0.004 \pm 0.003$<br>$t(302) = -1.36, p = 0.17$ |
|  | Exp. 4 | Exp. 5 | Exp. 6 |
| Intercept | $0.634 \pm 0.027$<br>$t(261) = \mathbf{23.06}, p < 0.001$ | $0.649 \pm 0.025$<br>$t(235) = \mathbf{26.30}, p < 0.001$ | $0.573 \pm 0.017$<br>$t(488) = \mathbf{34.30}, p < 0.001$ |
| Lag 2 | $0.010 \pm 0.005$<br>$t(261) = \mathbf{2.00}, p = 0.046$ | $0.012 \pm 0.004$<br>$t(235) = \mathbf{3.26}, p = 0.001$ | $0.007 \pm 0.002$<br>$t(488) = \mathbf{4.83}, p < 0.001$ |
| Lag 1 | $0.001 \pm 0.005$<br>$t(261) = 0.30, p = 0.78$ | $-0.005 \pm 0.004$<br>$t(235) = -1.48, p = 0.14$ | $0.007 \pm 0.002$<br>$t(488) = \mathbf{3.56}, p < 0.001$ |

#### Items repeated multiple times are retrieved faster, but the effect of recency is consistent across repetitions

Figure 5 shows median subject response times as a function of log lag and number of repetitions. Subjects were faster for items repeated more than once, but the effect of recency was consistent across repetitions. A linear mixed effects regression on median response times was performed treating subject as a random effect and log lag, number of repetitions, and the interaction of lag and repetition as fixed effects (Table 7). The same pattern of results emerged for all six experiments, there were significant main effects of log lag and repetition, but their interaction was not significant. That is, while participants were faster to respond the more times an item was presented, and were slower for less recent items, the delay to access those less recent items remained consistent regardless of how many times an item was presented. Taken together, these results indicate that the recency effect does not appear to be the result of better fidelity or improved encoding for more recent items, but rather arises at the comparison stage.

**Table 7:** Slope and intercept values from a linear mixed effects regression of lag and number of repetitions on median response times. All tests were performed on the base 2 logarithm of lag. Bold t-scores indicate significance at the *p* < 0.05 level. Across all six experiments, response times decreased with number of repetitions, but the effect of lag was the same regardless of number of repetitions.

| Median | Exp. 1 | Exp. 2 | Exp. 3 |
| --- | --- | --- | --- |
| Intercept | $0.934 \pm 0.042$<br>$t(179) = \mathbf{22.44}, p < 0.001$ | $0.602 \pm 0.012$<br>$t(182) = \mathbf{50.35}, p < 0.001$ | $0.674 \pm 0.020$<br>$t(187) = \mathbf{33.34}, p < 0.001$ |
| Slope | $0.013 \pm 0.004$<br>$t(179) = \mathbf{3.47}, p < 0.001$ | $0.013 \pm 0.002$<br>$t(182) = \mathbf{5.48}, p < 0.001$ | $0.012 \pm 0.003$<br>$t(187) = \mathbf{4.56}, p < 0.001$ |
| Repetition | $-0.069 \pm 0.023$<br>$t(179) = \mathbf{-2.96}, p = 0.004$ | $-0.067 \pm 0.009$<br>$t(182) = \mathbf{-7.52}, p < 0.001$ | $-0.080 \pm 0.009$<br>$t(187) = \mathbf{-8.52}, p < 0.001$ |
| Rep X Slope | $0.004 \pm 0.005$<br>$t(179) = 0.84, p = 0.40$ | $-0.004 \pm 0.003$<br>$t(182) = -1.13, p = 0.26$ | $0.002 \pm 0.004$<br>$t(187) = 0.52, p = 0.60$ |
|  | Exp. 4 | Exp. 5 | Exp. 6 |
| Intercept | $0.670 \pm 0.020$<br>$t(162) = \mathbf{34.06}, p < 0.001$ | $0.671 \pm 0.016$<br>$t(147) = \mathbf{42.30}, p < 0.001$ | $0.686 \pm 0.016$<br>$t(487) = \mathbf{42.45}, p < 0.001$ |
| Slope | $0.024 \pm 0.004$<br>$t(162) = \mathbf{5.87}, p < 0.001$ | $0.007 \pm 0.002$<br>$t(147) = \mathbf{3.03}, p = 0.003$ | $0.009 \pm 0.003$<br>$t(487) = \mathbf{3.36}, p < 0.001$ |
| Repetition | $-0.044 \pm 0.016$<br>$t(162) = \mathbf{-2.82}, p = 0.006$ | $-0.048 \pm 0.009$<br>$t(147) = \mathbf{-5.33}, p < 0.001$ | $-0.039 \pm 0.003$<br>$t(487) = \mathbf{-13.89}, p < 0.001$ |
| Rep X Slope | $-0.007 \pm 0.006$<br>$t(263) = -1.15, p = 0.25$ | $0.004 \pm 0.003$<br>$t(147) = 1.15, p = 0.25$ | $0.001 \pm 0.001$<br>$t(487) = 0.88, p = 0.38$ |

**Figure 5.**
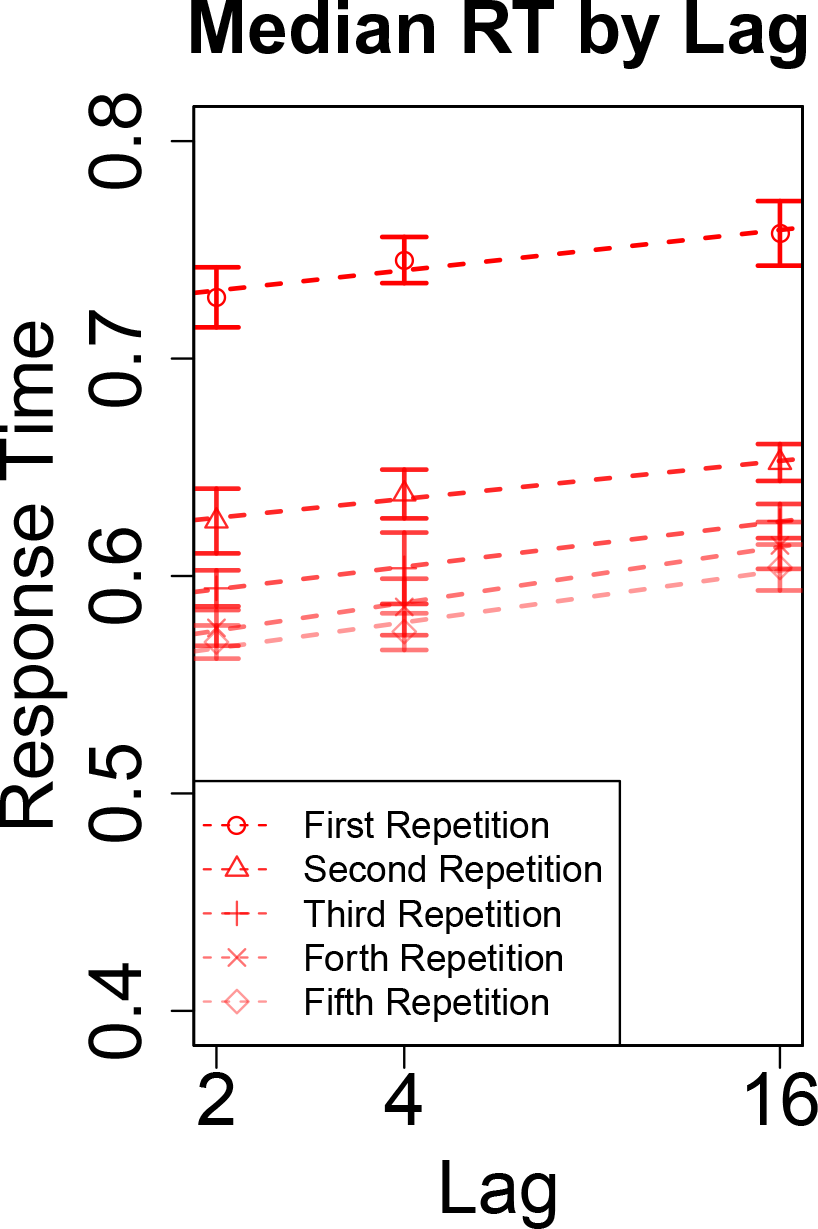
The effect of recency on response times was the same regardless of repetitions. Median response time as a function of the most recent lag and number of repetitions. To the extent the lines are parallel, it means that the effect of recency on median RT’s was the same for across repetitions. Analyses reported in the text demonstrated that only the most recent lag affected the RT of old probes repeated twice.

#### The time that items repeated multiple times become available depends on its recency

As demonstrated via Lag Modulation Factor, for items repeated once, the time an item becomes available depends on its recency (Figure 3). Although experiments 1-5 do not have enough trials to calculate a reliable Lag Modulation Factor for items repeated multiple times (9 trials per lag and subject), we can perform this on the data from experiment 6 (36 trials per lag and subject). We performed a mixed effect linear regression of Lag Modulation Factor onto quantile, treating subject as a random effect and quantile as a fixed effect. We found that Lag Modulation Factor was significantly different from zero at the intercept, 9.82 ± 1.11, *t*(244) = 8.87, *p* < 0.001, but did not significantly change as a function of quantile 1.29 ± 1.85, *t*(244) = 0.70, *p* = 0.49. That is, while there was an effect of recency for items repeated multiple times, for even the fastest responses, the effect of recency did not change throughout the distribution of responses. Taken together, these results offer strong support for the hypothesis that the recency effect is caused by the recovery of a temporal context, where the time to retrieve a context depends on its recency.

## General Discussion

It has long been known that response times in a continuous recognition experiment increase with the lag to the probe. If memory search requires accessing a timeline to find the appropriate memory, then the RT increase could be associated with a shift in the distribution. These six experiments showed that lag consistently shifted RT distributions. Consistent with this account, RT’s to second repetitions depended only on the most recent lag, as if the search terminated upon finding the first match. Although RTs were faster to second repetitions overall, the effect of the most recent lag on second repetition RTs was the same as the effect of lag on first presentation RTs (Figure 5). Consistent with earlier findings (e.g., Hockley, 1982), our results showed a sublinear shift with lag. This paper’s results are roughly consistent with a logarithmic increase in RT as a function of lag; each doubling of lag resulted in a shift of approximately 16-26 ms in the RT distribution.

At first glance, the results of this study are consistent with sequentially accessing a logarithmically-compressed timeline. There is extensive evidence for self-terminating serial search models in short-term memory tasks (Hacker, 1980; Hockley, 1984; McElree & Dosher, 1993; Sternberg, 2016). There is also evidence consistent with scanning along a timeline in judgement of recency tasks (Hintzman, 2010; Tiganj, Singh, Esfahani, & Howard, 2021). To be clear however, we are not suggesting that what we see here is scanning. The effect of doubling lag in our experiments resulted in an increase in RT of 16-25 ms, much faster than the increases seen in tasks believed to show scanning.

Most previous item recognition studies however have found evidence for parallel memory access, not sequential, in study-test recognition (Nosofsky, Little, Donkin, & Fific, 2011; Ratcliff & Murdock, 1976; McElree & Dosher, 1989; Hockley, 1984; Nosofsky, Cox, Cao, & Shiffrin, 2014). Several potentially important methodological differences may account for the discrepancy between those studies and this paper’s results. This experiment used continuous recognition rather than the study-test procedure (Nosofsky et al., 2011; Ratcliff & Murdock, 1976; McElree & Dosher, 1989; Hockley, 1984; Nosofsky et al., 2014). In addition, more recent lags were tested more frequently than more remote lags, new probes occurred far more often than repeated probes, and repeated probes could not appear in adjacent trials. The question of which combination of these methodological differences accounts for evidence supporting a timeline is a significant one that merits further investigation. It is worth noting that there is no reason in principle that a compressed timeline could not be accessed in parallel (Howard et al., 2015), whereas it is not clear how (or why) a composite representation could allow for changes in the time at which items become available. Notably, if a logarithmically-compressed timeline is accessed in parallel, the recency effect for the strength of match would fall off like a power law (Howard et al., 2015), much like the change Donkin and Nosofsky (2012) observed experimentally in drift rate.

Logarithmic compression is ubiquitous in psychology. Behaviorally, one way this manifests is via the Weber-Fechner law, the finding that the perceived intensity of stimulus changes with the log of its actual intensity (Fechner, 1860/1912). It has also been well established that compression is seen in neural populations. For example, the representation of visual space is logarithmically compressed as a function of distance from the fovea (E. L. Schwartz, 1980; Van Essen, Newsome, & Maunsell, 1984). Compression is also seen in time cells, neurons with temporal receptive fields which consistently fire at a specific time during a delay. Time cells show compression in that more cells have receptive fields earlier in the delay, and later firing cells have larger receptive fields (Tiganj, Cromer, Roy, Miller, & Howard, 2018; Cruzado, Tiganj, Brincat, Miller, & Howard, 2020). A recent analysis of time cells in rodent hippocampus, an area critical to episodic memory function, demonstrated that time cell compression is in fact logarithmic (Cao et al., 2021), such that, their distance in neural space goes up with log lag.

Episodic recollection is believed to be the result of occurs following the recovering a previous spatiotemporal context. This recovery results in behavioral changes, such as the contiguity effect. Previous recognition studies have found evidence that following episodic recollection, there is a boost in performance for items presented close in time to the just recovered item (G. Schwartz et al., 2005; Folkerts et al., 2018). In addition to behavioral effects, episodic recollection is accompanied by a neural contiguity effect as well (Folkerts et al., 2018), such that the similarity of the population at test is most similar to when the just recollected item was first presented, but also shows a high similarity to other items presented around the same time at study. The present datasets do not allow for us to measure a contiguity effect, as adjacent trials did not contain repetitions and we do not have a measure to separate “episodic recollection” trials from more familiarity base recognition. Despite this, our results are consistent with the hypothesis that the time to recover a temporal context depends on its recency in log space.

## Acknowledgments

The authors gratefully acknowledge useful discussions with Rui Cao, Zahra Esfahani, Zoran Tiganj, Anna Schapiro, Craig Stark, Jay McClelland, Rich Shiffrin, Rob Nosofsky, and Brandon Turner.

## Author Notes

This research was supported by NSF IIS 1631460 (IB, IS, MWH) and PHY 1444389 (MWH, IS) and a Google award (AO). All data and code used to perform analyses have been made publicly available on Github and can be accessed at https://github.com/tcnlab/ConRec. An earlier version of this manuscript containing the first three experiments was previously published on bioRxiv. It can be found at this link: https://www.biorxiv.org/content/10.1101/101295v1.

